# Single-nucleus analysis reveals renal STAT1/CEBPB signaling as a modifiable relay of proinflammatory FGF23 effects in diabetic kidney disease

**DOI:** 10.64898/2026.11.10.765607

**Authors:** Matthias B. Moor, Cheng Yang, Simge Sengul, Mikhail Burmakin, Jaakko Patrakka, Ping Chen, Hannes Olauson

## Abstract

**Background:** Circulating fibroblast growth factor 23 (FGF23) rises dramatically in chronic kidney disease, particularly in diabetic kidney disease (DKD). At high concentrations, and with declining Klotho, FGF23 has been linked to renal inflammation. The cell types and transcriptional programs mediating these off-target actions remain undefined. We asked whether the diseased kidney responds to FGF23 differently from the healthy kidney, and whether any disease-specific response could be targeted.

**Methods:** Male and female DBA/2J mice received streptozotocin or vehicle to induce DKD. On day 56, mice received a single dose intravenous injection of FGF23 (0.3 µg) or vehicle, with tissue collected 1 h later for single-nucleus RNA sequencing, differential expression and regulon (inferred transcription factor activity) analysis. Candidate pathways were tested in precision-cut kidney slices using the CEBPB inhibitor ST-101.

**Results:** Streptozotocin-treated male mice developed early DKD with a 6-fold increase in urinary albumin/creatinine ratio and 50% higher blood urea nitrogen with preserved glomerular filtration rate. All major renal cell types were annotated. Canonical FGF23 signaling was intact in disease, with segment-specific target induction conserved in both conditions (e.g. Cyp24a1 in PT-S1 and DCT1). Superimposed on this were disease-restricted responses, including *Wsb1* in DCT1 and *Pdk4* in PT-S2, PST, DCT1 and DCT2. Regulon analysis revealed DKD-specific induction of CEBPB regulon activity by FGF23 across all analyzed tubular segments, and of STAT1 in several segments. Pre-treatment with a CEBPB inhibitor suppressed CEBPB activity, downstream NF-κB and pro-inflammatory gene sets, and immune-like cell abundance in diseased slices.

**Conclusion:** In the diabetic kidney, FGF23 engages a STAT1-CEBPB-NF-κB axis absent from healthy kidney and extending beyond its canonical Klotho-dependent target segments, identifying CEBPB as a potential treatment target for FGF23’s pathological inflammatory actions.

## Introduction

One in three adults with diabetes mellitus have diabetic kidney disease (DKD), the most frequent cause of chronic kidney disease (CKD) globally (1). During the progression of DKD and CKD in general, the number of functional nephrons and renal expression of the protein Klotho declines, and the capacity of the kidney to excrete phosphate via the urine decreases (2). Consequently, many patients with DKD experience deteriorating kidney function and eventually develop hyperphosphatemia, soft-tissue calcifications and accelerated cardiovascular disease. Due to low-grade chronic inflammation in CKD, but also as a compensation mechanism to overcome the reduced phosphate excretion in advanced CKD, serum concentrations of the phosphaturic hormone Fibroblast Growth Factor 23 (FGF23) increase dramatically. In diabetes mellitus, either diminished levels or resistance to insulin contribute to rise in FGF23, as insulin cannot adequately repress *Fgf23* expression (3,4). Consequently, the pathophysiology of FGF23 excess is of particular importance in diabetes mellitus and predicts onset and progression of DKD (5,6), in addition to its well-established association with adverse cardiovascular outcomes in CKD in general (7).

In healthy kidneys, FGF23 acts on a receptor complex of Klotho and FGFR to reduce reabsorption of phosphate and increase urinary phosphate excretion (8). In advanced CKD, the combination of extremely elevated FGF23 concentrations and tissue-level resistance in conventional target cells with diminishing Klotho co-receptor abundance may in combination lead to adverse off-target effects and activation of non-canonical signaling pathways in the kidney (9). Aberrant FGF23 signaling in kidney disease models has recently been linked to renal chemokine secretion and immune cell recruitment through crosstalk between FGF23-driven MAPK signaling and NF-κB-dependent TNF-receptor activation, along with segment-specific inflammatory transcriptional responses (10). We recently established that substantial renal chemokine secretion and immune cell increase are consequences of aberrant FGF23 signaling in kidney disease models (11). The exact cell specificity and cell-specific pathways involved in these renal FGF23 off-target effects during CKD remain however a gap in knowledge. As such a proinflammatory renal environment is a key driver of later kidney fibrosis, it is pivotal to better understand the underlying mechanisms and identify druggable candidate pathways to alleviate aberrant renal cell signaling of FGF23.

Here, we aimed to improve our understanding of the cell-specific dynamics of renal FGF23 signaling in kidneys affected by DKD, in direct comparison with the corresponding changes in kidneys of healthy mice. We specifically hypothesized that FGF23 induces abnormal (“adverse”) signaling in DKD in mice, resulting from a combination of signaling in atypical, normally unresponsive cell types, and/or atypical transcriptional responses that are absent in healthy kidneys experiencing FGF23 signaling. We used an in-depth single-nucleus RNAseq screen combined with regulon analysis to identify the major drivers of FGF23 toxicity in a DKD model in mice, and we mechanistically validated the findings using pharmacological inhibition and precision-cut kidney slices.

## Methods

### 1. In vivo animal experiments

#### 1.1 Ethical approval and husbandry

All animal experiments were carried out at Karolinska Institutet following the protocols approved by the Ethical Committee on Research Animal Care (Linköping Animal Experimentation Ethics Committee; DNR 17707-2024). DBA/2J mice were purchased from Charles River. Mice were housed in individually ventilated cages under a 12h light, 12h dark cycle with unlimited access to standard chow (Lantmannen) and water. Temperature was constant at 20 ± 2°C and humidity was 50 ± 5%. For all animal experiments, euthanasia methods were terminal exsanguination followed by cervical dislocation under anesthesia with isoflurane USP (Baxter), followed by organ collection.

#### 1.2 Induction and monitoring of DKD model in mice

For induction of experimental diabetes, 8 week-old mice were used. Streptozocin (Sigma #S0130) was applied dissolved in 0.1M sodium citrate buffer at pH 4.5. Males were intraperitoneally injected with 40 mg/kg Streptozocin or citrate buffer, and females with 50 mg/kg Streptozocin or citrate buffer for five consecutive days, to account for the relative lower STZ sensitivity of females (12,13). Blood and urine samples were obtained at baseline, at day 15 and weekly thereafter for biochemical analyses.

#### 1.3 Hormone injections

Recombinant murine FGF23 was purchased from PeproTech (distributed via ThermoFisher, #450-55). On day 56 after induction of DKD, FGF23 hormone was intravenously administered using a protocol adapted from earlier studies (11,14) and a pilot dose-finding study. Mice of both sexes with and without DKD were randomly allocated to receive a single intravenous injection of 0.3 μg of FGF23 dissolved in PBS-BSA 0.1% (Sigma) or PBSA-BSA 0.1% vehicle. One hour after FGF23 administration, spot urines were collected and mice were euthanized for blood and organ collection.

### 2. Serum and urine biochemistry

Biochemical analyses were made by commercial kits according to manufacturer instructions. We measured urinary albumin concentration using the Albuwell M ELISA kit (#1011, Ethos), blood urea nitrogen (BUN) using the BUN Colorimetric Detection Kit (EIABUN, Thermo Fisher), creatinine by a colorimetric detection kit (AB65340-1001, Abcam), glucose by FreeStyle Lite blood glucose monitor system (Abbott) and phosphate by a colorimetric kit (Cat. MAK-030, Merck). Murine FGF23 was quantified using an ELISA kit for intact FGF23 (#60–6800, Immutopics, Inc.).

### 3. Ex vivo precision-cut kidney slices experiments

#### 3.1 PCKS preparation

PCKS were established from a separate group of male DBA/2J mice undergoing the same protocol with Streptozocin-induced DKD and controls, using a previously established protocol (11,15). A Compresstome® VF-310-0Z Vibratome (Precisionary Instruments) was used after preparation with William’s E Medium GlutaMAX-I buffer (WME solution, Thermo Fisher) with 25 mM D-glucose (Merck). Pairs of 300 µm kidney slices were made per mouse. The slices were transferred to 12-well plates and put into 1.5 mL of WME solution at 37°C and 95% oxygen for a total experiment duration of 2h.

#### 3.2 Ex vivo treatments

Experimental treatments were given to three groups of two PCKS per animal for *ex vivo* treatment with either PBS-BSA 0.1% vehicle for 1h, recombinant murine FGF23 at 200 ng/mL for 1h, or recombinant murine FGF23 at 200 ng/mL for 1h under a co-treatment of CEBPB inhibitor ST-101 / Lucicebtide (#PC-49350, ProbeChem) 20 µmol/L during the entire 2h experiment, a protocol adapted from (16). Slices were then collected, snap-frozen and stored at -80°C until further processing.

### 4. RNA and nuclei isolation

#### 4.1 Total RNA isolation

Tissue homogenization of approximately 30 mg of renal *pars intermedia* were performed using a TissueLyser in 600 μL RLT buffer with DTT. RNA extraction steps were then followed using the RNeasy Mini Kit (Qiagen) according to manufacturer instructions.

#### 4.2 Single-nuclei isolation

Nuclei were isolated from a piece of snap-frozen *pars intermedia* kidney cortex that weighed 30-50mg, using a method adapted from Legouis et al. (17). Tissue was minced by scissors and thawed in 500 uL of ice-cold nLB containing RNAse inhibitors. The tissue was then homogenized by 10-20 gentle rotations of an RNase/DNase-free mini-brush (Milan #1005-39), followed by lysis and filtration through 70 and 30μm cell strainers. Nuclei were then washed twice in NWSB and spun down, followed by filtration through a 35μm cell strainer, and staining by trypan blue for manual counting by hemocytometer. 20.000 nuclei per sample were then suspended at 1500 nuclei/μL in PBS containing 0.8% BSA, 0.5% RNase inhibitor and 10% DMSO.

### 5. RNA analysis workflows

#### 5.1 qPCR

Renal RNA was processed for cDNA synthesis using the iScript cDNA Synthesis Kit (Bio-Rad). Next, a CFX96Real-Time PCR Detection System and iQ SYBR Green Supermix (Bio-Rad) were used for detection of relative gene expression by the 2^− ΔΔCT^ method. Primers were mm*Gapdh*-F: 5′tgttcctacccccaatgtgt3′, mm*Gapdh*-R: 5′tgtgagggagatgctcagtg3′, mm*Egr1*-F: 5’agccgagcgaacaaccctat3’ mm*Egr1*-R: 5’cgccttctcattattcagagcg3’.

#### 5.2 Bulk RNAseq

Quality of m RNA was checked with the Agilent TapeStation. Illumina TruSeq Stranded mRNA ligation was used for library preparation. A standard protocol of cDNA synthesis, ligation of adapters, and amplification of indexed libraries was used. Yield and quality of the libraries were analyzed using Qubit by Thermo Fisher and the Agilent TapeStation. The cDNA libraries were normalized and combined. Pools were sequenced on an Illumina NextSeq 2000 platform with a P3 100 sequencing run generating 58 bp paired-end reads.

#### 5.3 Single-nucleus RNAseq

Integrity and concentration of the nuclei from 16 samples was assessed using a Luna FL cell counter (Logos Biosystems). Next, the nuclei were loaded on a 10x Genomics GEM-X loading chip, aiming for a target recovery of 15.000 nuclei per sample. The droplet encapsulating device was a 10x Genomics Chromium iX. Next, sequencing libraries were generated according to manufacturers’ instructions. The libraries were then pooled in sets of eight samples, and sequenced on an Illumina NovaSeqX 25B flow cell, with eight multiplexed samples per lane, with aim to read 375m reads per sample, or 25.000 reads per nucleus. The resulting data was then demultiplexed and preprocssed using the 10x Genomics Cellranger software (Version 8.0) and the mouse GRC38 (mm10) genomic build, provided as a prebuild reference transcriptome by 10x Genomics.

### 6. Bioinformatics and statistical analyses

#### 6.1 Biochemical data analysis

Serum and urine biochemistry data were analyzed using two-way analysis of variance (ANOVA) in GraphPad Prism 8.0, assuming normally distributed data. Disease status (DKD vs. healthy) and hormone treatment (FGF23 vs. vehicle) were included as the two factors. qPCR data were analyzed by ANOVA.

#### 6.2 Bulk RNAseq data analysis

Bulk RNAseq data were demultiplexed using bcl2fastq (v2.20.0) and sequencing reads were aligned to the mouse reference genome (GRCm39) using STAR (v2.7.9a). Aligned reads were assigned to coding regions of biotypes from Ensembl Mus_musculus.GRCm39.113.gtf using featureCounts (v1.5.1). Count data were normalized using Trimmed Mean of M-values. Differential gene expression were analyzed using limma (v3.64.3). Four paired comparisons were performed to access (i) the FGF23 effect in PCKS tissues from healthy mice, (ii) the CEBPBi effect in FGF23-treated PCKS from healthy mice, (iii) the FGF23 effect in PCKS from mice with DKD, and (iv) the CEBPBi effect in FGF23-treated PCKS from DKD mice separately. Gene set enrichment analysis (GSEA) was performed on logFC-ranked gene lists using clusterProfiler (v4.16.0) package with the Molecular Signatures Database (MSigDB) collection of gene sets. In addition, MSigDB gene set GERY_CEBP targets (MM1155) was separately visualized using the gseaplot function. For estimation of immune-like cell fractions, RNAseq data underwent bulk deconvolution against single-cell RNAseq reference data from healthy mice (GSE107585) using Seurat 5.0.3 and R/Bisque v.1.0.5 as previously reported (11,18,19). Immune-like cell fractions were visualized and compared using nonparametric repeated-measures ANOVA (Friedman test) in GraphPad Prism 8.0. Bulk transcription factor activity was inferred from t statistic of limma contrasts using R/Dorothea 1.20.0 and R/decoupleR 2.14.0.

#### 6.3 Single-nucleus data analysis

##### 6.3.1 Data preprocessing and quality control

Single-nucleus RNA-seq data of all 16 samples were merged and analyzed in Python using the Scanpy framework (v1.10.2), retaining sample identity, disease status (DKD or healthy), and treatment condition (FGF23 or vehicle) as metadata. Quality control was performed sequentially using sample-specific and global filtering criteria. First, thresholds were set for each sample after inspecting the distribution of each quality metric, including total UMI count, the number of detected genes, and the fractions of counts mapping to mitochondrial, ribosomal, and hemoglobin genes. Nuclei with low gene complexity (fewer than 350 to 850 detected genes, depending on the sample) or elevated mitochondrial read fraction (above sample-specific threshold ranging from 0.1% to 5%) were excluded. In addition, putative low-quality nuclei with disproportionately high UMI counts (> 2,500) but low gene detection (typically < 1,000 genes) were removed. After sample-level filtering, potential doublets were identified using Scrublet (v0.2.3) and removed with a doublet score threshold of 0.25. Following inspection of sample-level quality metric distributions, a final global quality-control step was applied to remove remaining outliers. Nuclei with mitochondrial read fractions >1%, hemoglobin gene fractions >5%, or total UMI counts or detected gene counts above the 99th percentile were excluded. Raw count matrices were preserved in a dedicated *counts* layer prior to normalization.

##### 6.3.2 Cell type annotation

Gene expression counts were normalized to a total of 10,000 counts per nucleus, followed by log1p transformation in Scanpy. The 3,000 most highly variable genes (HVGs) were selected using flavor=‘cell_ranger’. Principal component analysis (PCA) was performed on HVGs using the ARPACK sparse solver, retaining 50 principal components (PCs). A k-nearest-neighbor (kNN) graph was constructed using the top PCs and 15 nearest neighbors. The data were visualized in two dimensions using uniform manifold approximation and projection (UMAP). Unsupervised clustering was performed using the Leiden algorithm at a resolution of 0.5, with a fixed random seed of 42 for clustering and 34 for UMAP. Marker genes for each cluster were identified using the *t*-test implemented in Scanpy. Cluster identity was assessed using the marker_gene_overlap function in Scanpy against a curated panel of canonical kidney cell-type markers.

Initial cell type annotation was performed using Leiden clustering at a resolution of 0.5, yielding 18 clusters. Cell identities were manually assigned based on the expression of canonical kidney cell-type markers, including proximal convoluted tubule S1 (PCT/PT S1: *Slc5a12, Slc34a1, Pdzk1*), proximal straight tubule (PST: *Slc22a30, Slc13a3, Cyp4b1*), thick ascending limb (TAL: *Slc12a1, Umod, Cldn19*), distal convoluted tubule (DCT: *Slc12a3, Calb1, Pvalb*), connecting tubule/principal cells (CNT/PC: *Aqp2, Hsd11b2, Fxyd4*), intercalated cells (IC: *Atp6v1g3, Atp6v0d2, Car2*), podocytes (Podo: *Nphs1, Nphs2, Magi2*), endothelial cells (EC: *Pecam1, Cdh5, Plvap*), mesangial/myofibroblast-like cells (MLC: *Acta2, Cnn1, Pdgfrb*), parietal epithelial cells (PECs: *Cldn1, Dkk3, Wt1*), immune cells (*Ptprc, Cd68, Lyz2*), and macula densa (MD: *Slc8a1, Slit2*). Based on these markers, cell clusters were initially annotated as PCT 1, PCT 2, PCT 3, DCT 1, TAL, EC 1, DCT 2, PST, PC, MLC, IC, PECs, EC 2, Podo, Immune, MD.

Selected cell populations were further resolved by iterative sub-clustering at a Leiden resolution of 0.2, using a dedicated sub-clustering function applied to the neighbor graph of the full dataset. Intercalated cells were divided into type A (ICA: *Pam, Tmem117, Itpr2*) and type B (ICB: *Slc26a4, Slc4a9*) subtypes, along with a putative juxtaglomerular apparatus (JGA) population characterized by *Pbx1* and *Meis2* expression. Sub-clustering of parietal epithelial cells (PECs) revealing an injured proximal tubule cell sub-population (Inj.PT: *Havcr1, Vcam1, Dock10*). Endothelial cells were further resolved into glomerular/peritubular EC (*Ehd3, Emcn*) and a *Slc12a1*-expressing endothelial subset. To align with standard nomenclature, PCT 1 was renamed PT S1, while PCT 2 and PCT 3 were merged into PT S2. The final annotated dataset comprised 18 cell populations: PT S1, PT S2, PST, TAL, DCT 1, DCT 2, CNT, ICA, ICB, JGA, Podo, EC 1, EC 2, MLC, MD, PECs, Inj.PT, and Immune, visualized on a UMAP embedding (Figure 1). Dot plots of top marker genes per cell type were generated using Scanpy’s *dotplot* function with standard scaling per gene, expression cutoff of 0.99, and the plasma colormap.

**Figure 1.**
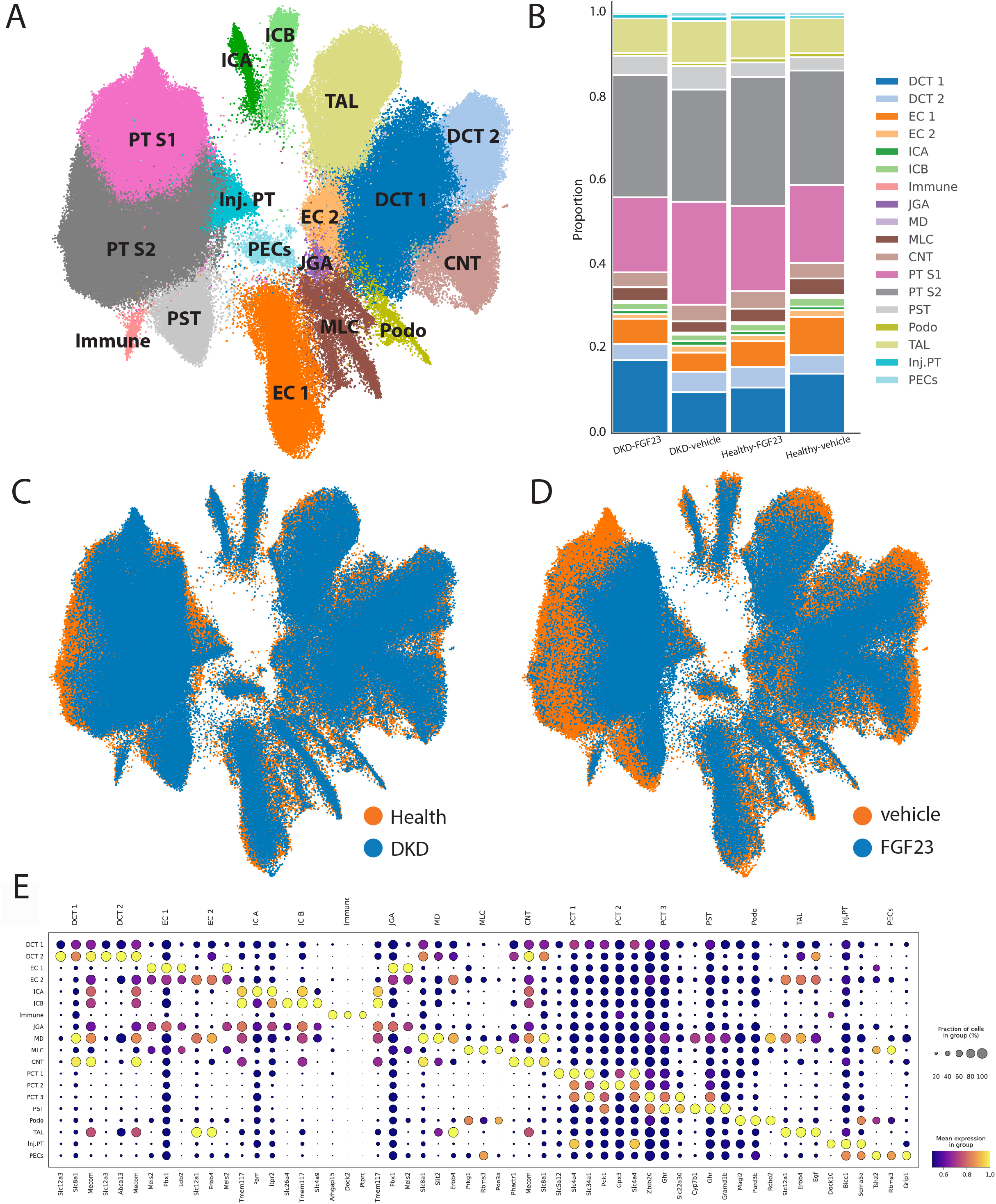
Annotation of the major nephron segments in healthy mice and mice with diabetic kidney disease (DKD). UMAP of 18 major kidney cell types (A) and cellular composition of the 4 experimental groups with and without DKD and FGF23 vs vehicle treatment (B). All groups provided cells to every cluster independent of disease status [DKD vs healthy] (C) or treatment status [FGF23 vs vehicle] (D). E shows top 4 transcripts per cluster.

##### 6.3.3 Differential gene expression and functional analysis

Differential gene expression analysis was performed in R (v4.4.1) using MAST (v1.30.0) to assess the effects of FGF23 treatment within each tubular cell population (PT S1, PT S2, PST, CNT, DCT 1, DCT 2, and Inj.PT) under healthy and diabetic kidney disease (DKD) conditions. For each cell, The cellular detection rate was calculated and included as a covariate to account for differences in gene detection. For each cell type and disease-status subset, MAST hurdle models were fitted with treatment (FGF23 vs. vehicle) as the primary variable of interest, with vehicle set as the reference. The treatment effect was assessed using a likelihood ratio test. Subsets lacking representation of both treatment groups were excluded from the analysis. P values were adjusted for multiple testing using the Benjamini-Hochberg method, and significant differentially expressed genes were identified using adjusted P value < 0.05 and |log_2_FC| > 0.3.

To characterize similarities and differences in transcriptional responses to FGF23 between healthy and DKD conditions, DEGs identified under healthy and DKD conditions were compared within each cell type and classified as healthy-specific, DKD-specific, or common DEGs using custom R scripts. Gene symbols were converted to Entrez Gene identifiers using the bitr function against the org.Mm.eg.db annotation database (v3.19.1). DEG sets containing fewer than five genes with successfully mapped Entrez identifiers were excluded from subsequent enrichment analysis. Gene Ontology (GO) Biological Process and KEGG pathway over-representation analysis was performed separately for healthy-specific and DKD-specific DEGs for each cell type using clusterProfiler (v4.12.6).

##### 6.3.4 Regulon analysis

Gene regulatory network (GRN) inference was performed using the pySCENIC Singularity container (aertslab/pyscenic:0.12.1) following the standard pySCENIC workflow (20). The raw single-cell expression matrix was converted into a loom file and used as input for the pySCENIC pipeline. Co-expression modules between transcription factors and candidate target genes were identified with GRNBoost2, run three times to account for the stochastic nature of the algorithm. For each run, co-expression modules were pruned based on transcription factor binding motif enrichment using the mouse cisTarget database, and regulons were defined by retaining only target genes with supporting motif evidence. Transcription factor-target gene interactions identified in at least two of the three runs were retained and used for regulon activity scoring with AUCell. The resulting regulon activity score matrix and gene regulatory network were then used for downstream analyses. Cell-type specificity of regulons was then quantified using the Regulon Specificity Score (RSS) (21), computed with pySCENIC across groups defined by cell type, condition, and treatment.

## Results

### Sex-dependent induction of diabetic kidney disease in mice

We and others have previously reported that inflammatory responses are induced by FGF23 excess in using different disease models in mice (11,22,23). In the present study, we aimed to systematically decipher the renal cell-type-specific responses to FGF23 and distinguish conserved signaling mechanism from those associated with diabetic kidney disease (DKD). To this end, we used a classical model of Streptozotocin-induced diabetic kidney disease with sex-adapted doses in male and female mice.

First, we set out to characterize the DKD model and determine the optimal time point of experiments. Blood glucose analyses revealed an induction of elevated glycemia in both sexes from 15th day on which was more pronounced in males than females (Supplemental Fig. 1). Early signs of kidney damage are sensitively assessed by elevated urinary protein excretion, so we assessed the urinary albumin/creatinin ratio (ACR). ACR was elevated in males from day 21 on through day 56 but remained unchanged in most of females (Supplemental Fig. 1). Healthy males gained more weight than males with DKD. Renal retention parameters blood urea nitrogen (BUN) was increase in males with DKD but not in females, and serum creatinine concentrations were increased in mice with DKD of both sexes. With the expected glomerular hyperfiltration during early DKD, glomerular filtration rate (GFR) was not reduced in the groups with DKD (Supplemental Fig. 1).

Overall, male STZ-treated mice showed a robust presence of DKD after 8 weeks. As proteinuria, the classic clinical hallmark of DKD, was missing in females, we restricted the following analyses to males.

### Biochemical profiling of FGF23 effects

We used a pilot experiment for dose-finding of the minimum dose to induce a cell signaling effect of FGF23. This pilot experiment revealed a most robust induction of renal *Egr1* expression at 300ng of recombinant FGF23 (Supplemental Figure 2). Consequently, we intravenously treated male mice with and without DKD with 300ng of recombinant FGF23. While the focus of the present work was cellular signaling, we also assessed the effect of FGF23 on biochemical parameters. The short 1-hour treatments with FGF23 caused an increase in circulating iFGF23 (Supplemental Figure 2) and fractional excretion of phosphate (FE_Pi_) regardless of disease state (Supplemental Figure 2). In addition, DKD caused an overall increase in FE_Pi_ (Supplemental Figure 2), whereas phosphatemia remained comparable (Supplemental Figure 2).

**Figure 2.**
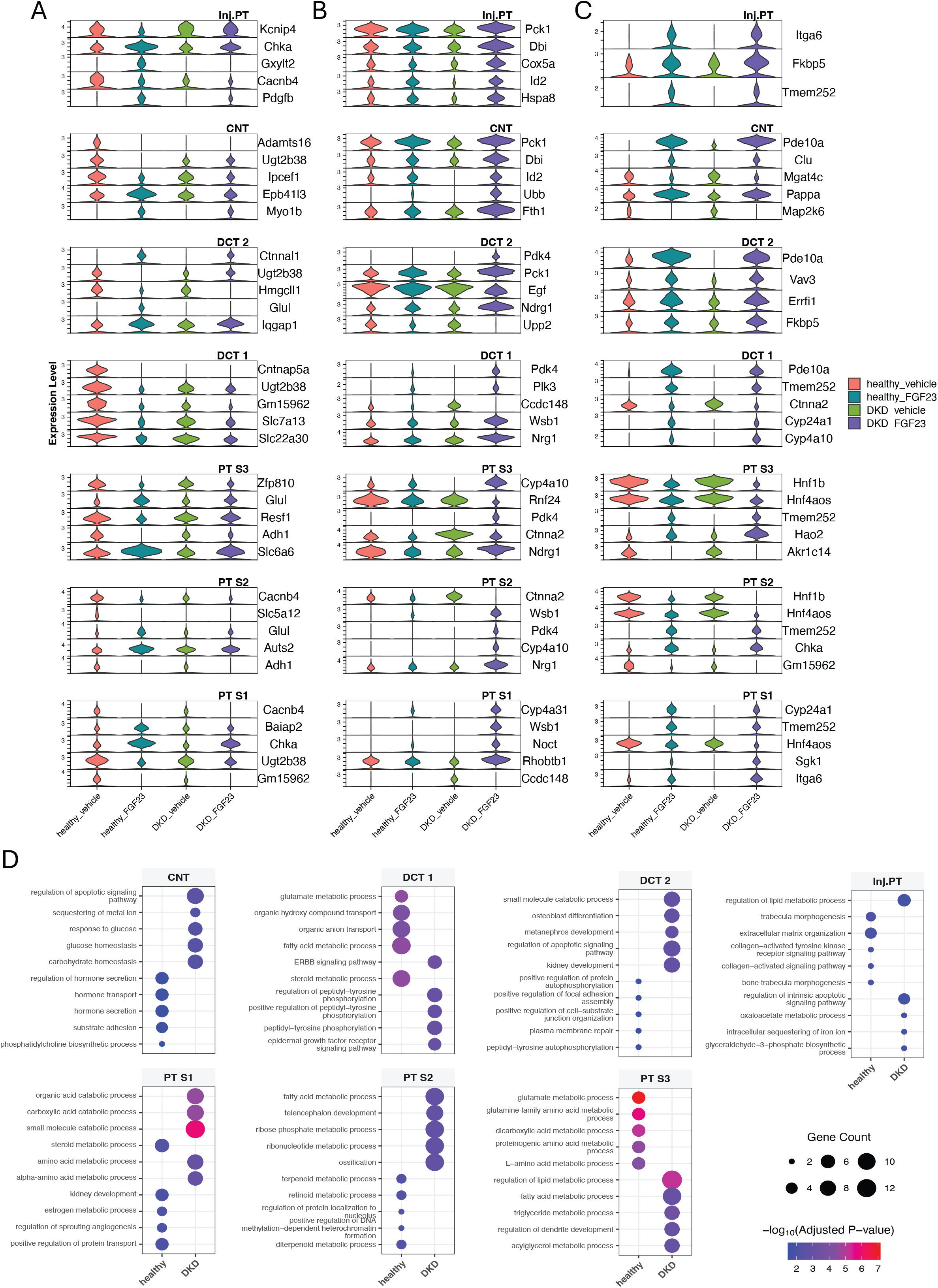
FGF23-driven transcriptional effects in the FGF23-responsive nephron segments shift during DKD. A shows healthy-specific and B shows DKD-specific transcriptional responses by displaying top 5 differentially expressed genes (DEG) in response to FGF23 in 4 proximal and 3 distal renal tubular nephron segments. C indicates the top 5 common FGF23-driven DEGs detected in both healthy and mice with DKD. D shows top gene ontology (GO) with top 5 healthy-specific and top 5 DKD-specific processes. PT, proximal tubule. PST, proximal straight tubule. CNT, connecting tubule. DCT, distal convoluted tubule. S1-2, segment 1-2.

### Annotation of the major nephron segments in healthy mice and mice with diabetic kidney disease (DKD)

We harvested the kidneys of 16 mice from all 4 groups for nuclei isolation. Approximately 20,000 extracted nuclei per sample were loaded on the 10x Chromium platform and sequenced. We used several quality control steps during data processing, leading to a total of 264.431 high-quality single nuclei which were annotated as 18 major kidney cell types (Figure 1A). The cellular composition of the four experimental groups was overall comparable in stacked barplots (Figure 1B). In addition, all experimental groups provided cells to every cluster, irrespective of disease status [DKD vs healthy] (Figure 1C) or treatment status [FGF23 vs vehicle] (Figure 1D). Overall, the average number of transcripts and distinct genes per nucleus was 3141 and 1615. All clusters displayed distinct transcript markers as expected (Figure 1E).

### Cell-type-specific transcriptional responses to FGF23 in healthy and diabetic kidneys

To investigate how diabetic kidney disease (DKD) influences renal responses to FGF23, we performed single-nucleus RNA sequencing (snRNA-seq) across four experimental groups comprising 16 mice. Following quality control, 264,431 high-quality nuclei were retained and classified into 18 distinct kidney cell populations, representing major nephron segments as well as vascular, stromal, and immune cells (Figure 1A). The overall cellular composition was broadly comparable across experimental groups (Figure 1B), with all identified populations represented in both healthy and DKD mice (Figure 1C) and across FGF23- and vehicle-treated conditions (Figure 1D). Each population exhibited distinct expression patterns of canonical marker genes, supporting their cell-type annotations (Figure 1E).

We next investigated the transcriptional responses to FGF23 within seven tubular cell populations, including four proximal and three distal tubular populations, under healthy and DKD conditions. Differential expression analysis revealed diverse transcriptional responses to FGF23 treatment between the two disease states. Among genes identified as differentially expressed under healthy conditions, FGF23 increased the expression of *Auts2*, encoding a chromatin-associated regulatory protein and *Glul* encoding glutamine synthetase in PT S2 cells (Figure 2A). In contrast, DKD-specific differentially expressed genes included *Pdk4*, encoding pyruvate dehydrogenase kinase 4, which was upregulated following FGF23 treatment in PT S2, PT S3, DCT 1, and DCT 2 cells (Figure 2B). Additional DKD-specific responses included increased expression of *Noct4* in PT S1, *Cyp4a10* in PT S3, and *Wsb1* in DCT 1. Despite these disease-associated differences, several FGF23 responses were shared between healthy and DKD conditions. Notably, FGF23 induced *Cyp24a1* expression in PT S1 and DCT 1 and *Sgk1* expression in PT S1, while reducing *Hnf1b* expression in PT S2 and PT S3 and *Map2k6* expression in CNT (Figure 2C).

To explore the biological processes associated with these transcriptional changes, we performed Gene Ontology (GO) over-representation analysis of FGF23-driven DEGs within each tubular population. Under healthy condition, FGF23-associated changes involved protein transport in PT-S1, dicarboxylic acid metabolism in PST, and steroid metabolic process in DCT1. In contrast, DKD-specific responses were enriched for amino acid and carboxylic acid metabolism in PT-S1, and glucose homeostasis and apoptosis-related signaling across several tubular populations. Notably, glyceraldehyde-3-phosphate biosynthesis was identified specifically in injured PT cells (Figure 2D).

Overall, these findings showed that FGF23-associated transcriptional responses vary across nephron segments and disease states. While several regulatory responses were shared between healthy and diabetic kidneys, DKD was associated with distinct transcriptional and metabolic pathway signatures, suggesting that the diabetic environment modifies the renal response to FGF23.

### Regulon activity analysis reveals aberrant FGF23-driven STAT1-CEBPB signaling in diabetic kidney disease

Given the divergent transcriptional responses to FGF23 observed between healthy and DKD kidneys, we next investigated whether these differences were associated with altered transcription factor regulatory programs. To this end, we employed regulon analysis by pySCENIC and computed regulon specificity score (RSS) for each experimental group in the FGF23-responsive nephron segments. In FGF23-treated DKD mice, CEBPB consistently ranked among the top regulons across the analyzed tubular populations (Figure 3A). STAT1 was also among the top-ranked regulons in several populations, together with JUN, JUND, and YBX1, although the latter were less consistently represented. Notably, CEBPB regulon specificity was markedly higher in FGF23-treated mice with DKD than in the other experimental groups (Figure 3B). A comparison with the top 5 regulons in the other experimental groups revealed a complete absence of CEBPB, and STAT1 was among the top regulons only in PT-S1 of vehicle-treated mice with DKD (Supplemental Figure 3).

**Figure 3.**
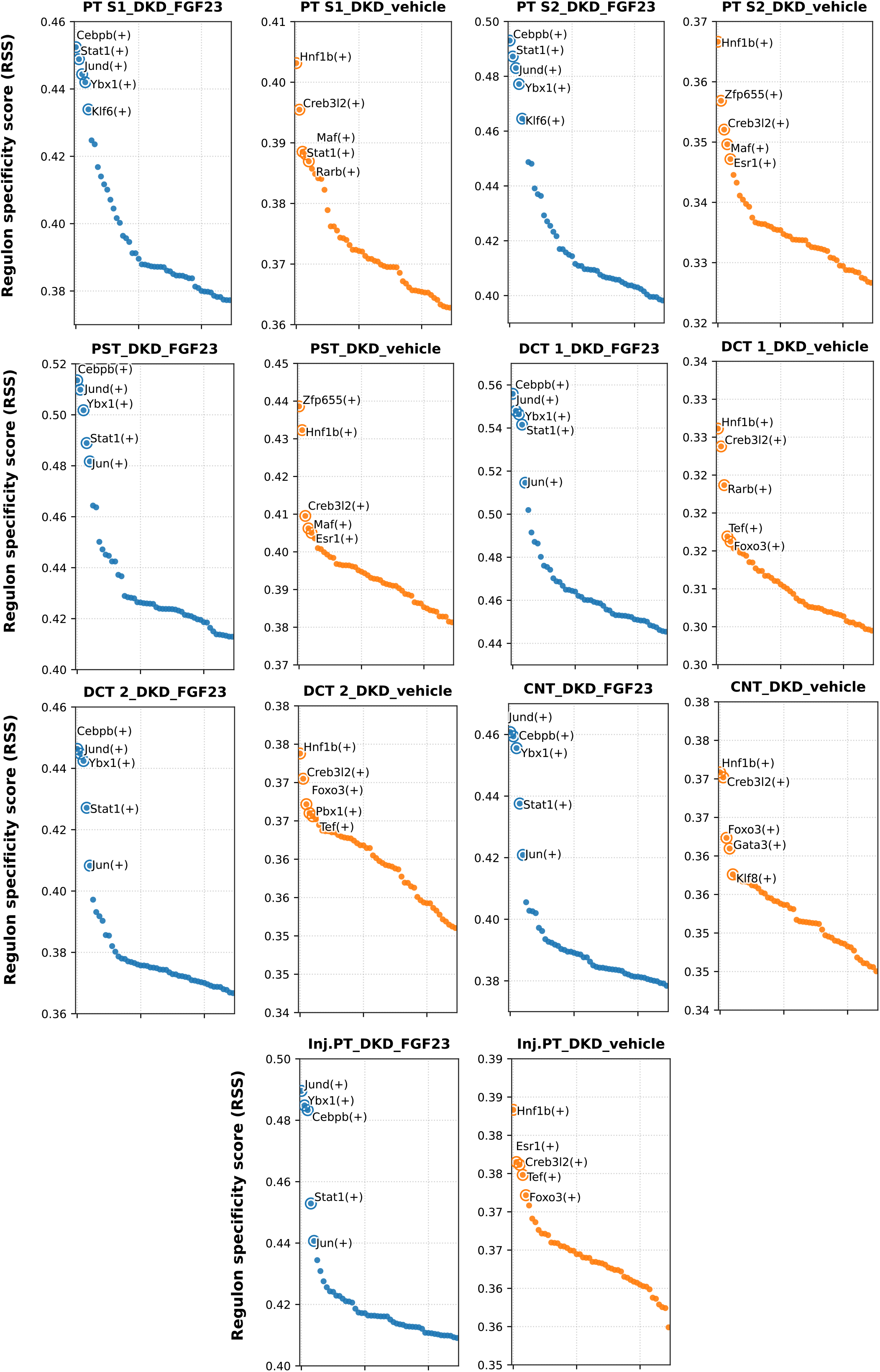
Regulon activity analysis reveals evidence of STAT1-CEBPB signaling upon FGF23 treatment in mice with DKD. pySCENIC analysis of FGF23-treated mice with DKD in 4 proximal and 3 distal nephron segments. PT, proximal tubule. PST, proximal straight tubule. CNT, connecting tubule. DCT, distal convoluted tubule. S1-2, segment 1-2. N=4 mice per group; RSS was computed on nuclei pooled within each experimental group.

However, in healthy mice, we found EGR1 among the top regulons upon FGF23 treatment in all of PT-S1, PT-S2, PST, DCT1, DCT2 and CNT as expected (10) (Supplemental Figure 3). FGF23 treatment was also associated with a shift in the predominant proximal tubular regulons, from HNF1B and HNF4G in vehicle-treated kidneys toward HNF4A in FGF23-treated kidneys. This finding was consistent with the reduced *Hnf1b* expression observed in PT S2 and PST following FGF23 treatment (Figure 2C). CEBPB was not among the top regulons in any of these other experimental groups.

In summary, we found a robust DKD-specific STAT1 and CEBPB regulon activity driven by FGF23 across nephron segments, whereas the physiological FGF23-driven EGR1 and HNF4A was predominant among the top regulons only in healthy kidney.

### Pharmacological CEBPB inhibition alleviates FGF23-driven proinflammatory action in diabetic kidney disease in an ex vivo model of precision-cut kidney slices

CEBPB is a proinflammatory transcription factor in diabetes (24–27). Based on our above findings, we thus hypothesized that CEBPB mediates some of the downstream proinflammatory effects induced by FGF23 in DKD. To test this, we harnessed an *ex vivo* set-up of precision-cut kidney slices (PCKS) from mice with and without DKD (Figure 4A). In this experiment, we treated PCKS with vehicle, with FGF23 or with a pre-treatment of CEBPB inhibitor ST-101 followed by FGF23. In GSEA analysis, we observed that the FGF23 treatment alone induced enrichment of CEBP target genes in PCKS, regardless of disease state (Figure 4B). Addition of the inhibitor ST-101 efficiently repressed the FGF23-induced CEBP targets enrichment (Figure 4B). We then specifically compared the top gene sets affected by ST-101 pre-treatment in PCKS: In DKD, ST-101 repressed pro-inflammatory pathways related to leukocyte chemotaxis, aggregation or migration (Figure 4C), but not in PCKS from healthy mice (Supplemental Figure 4). Further, NFKB1 was among the top TF activity scores repressed by ST-101 during FGF23 signaling in DKD (Figure 4D), but absent in PCKS of healthy mice (Supplemental Figure 5). Finally, the transcriptome-inferred relative abundance of cells reminiscent of immune cells was repressed by ST-101 in FGF23-treated PCKS of mice with DKD (Figure 4E). Overall, these data pointed towards a modifiable role of CEBPB in promoting pro-inflammatory transcriptional responses to FGF23 in DKD.

**Figure 4.**
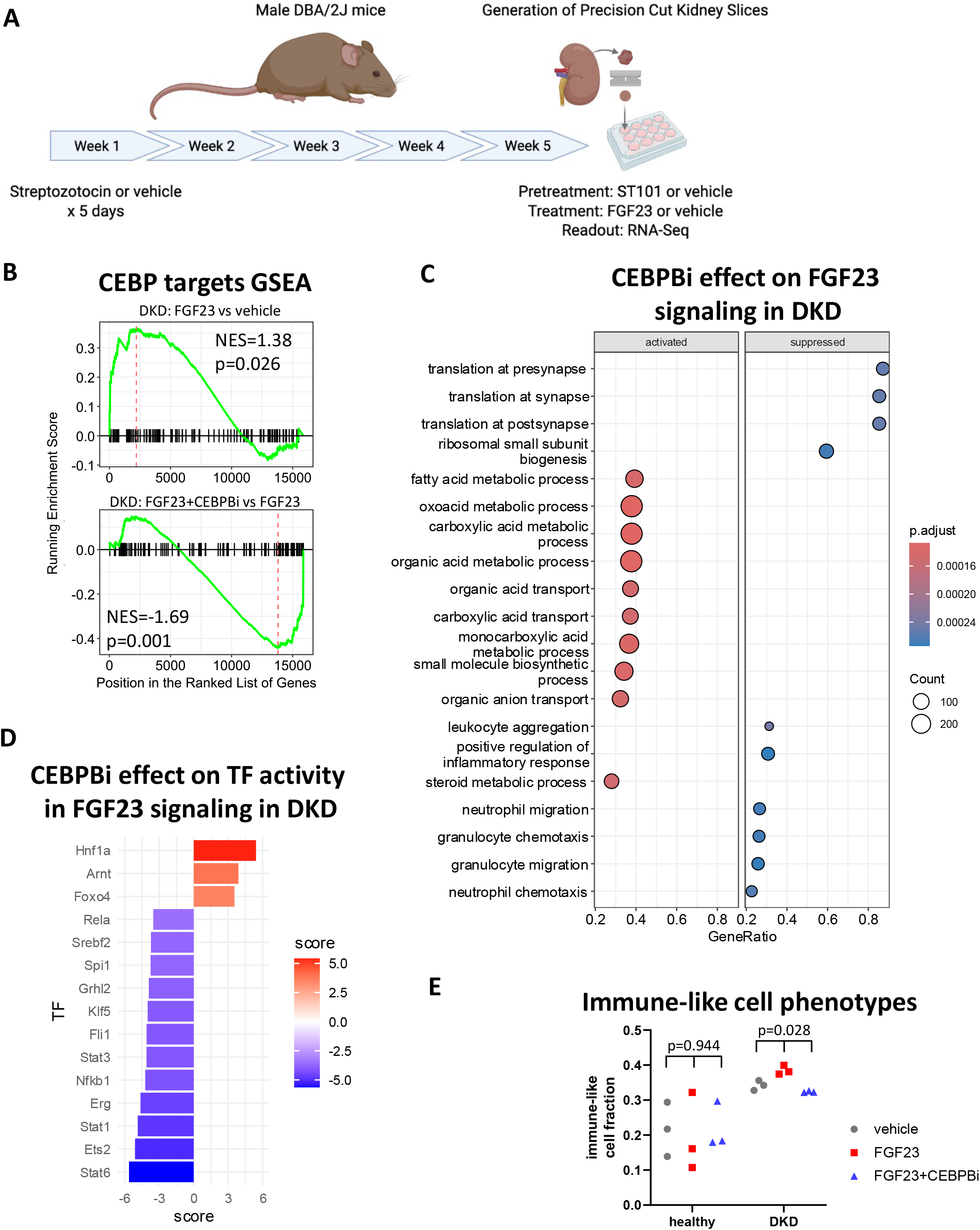
CEBPB inhibition alleviates FGF23-driven proinflammatory actions in an *ex vivo* model of DKD using precision-cut kidney slices (PCKS). A, Experimental algorithm, graphic made using a licensed version of bioRender. Effect of FGF23 vs vehicle (B, top) and FGF23 with/without CEBPB inhibitor ST-101 (B, bottom) on CEBP target gene set in PCKS of mice with DKD. C shows top enriched gene sets upon CEBPB inhibition during FGF23 signaling in PCKS of mice with DKD. D shows transcription factor (TF) activity modulated by CEBPB inhibition during FGF23 signaling in PCKS from mice with DKD. E displays transcriptome-inferred immune-like cell fractions in PCKS from healthy or mice with DKD as indicated. Paired contrasts, PCKS from N=3 healthy and 3 mice with DKD.

## Discussion

In this work, we aimed to cell-specifically decipher the pathological signaling events induced in the kidney by the hormone FGF23 at a disease state, with focus on a mouse model of early DKD treated acutely with a low dose of FGF23. We found that some normal signaling outcomes e.g. induction of *Cyp24a1* encoding a key vitamin D metabolizing enzyme, were preserved in DKD. However, our analyses highlight adverse signaling trajectories in response to FGF23 in DKD, including shifts of transcriptional signatures related to glucose metabolism and pro-apoptotic pathways in several nephron segments. Inferred TF activity analyses revealed DKD-specific STAT1-CEBPB activation upon FGF23 treatment along the nephron. Pharmacological inhibition of CEBPB alleviated NF-KB activation and proinflammatory gene expression during FGF23 signaling in an *ex vivo* model of PCKS from mice with DKD. These results reveal pathological cell signaling downstream of FGF23 in a preclinical model of diabetic kidney disease.

### Interpretation

We and others have previously reported pro-inflammatory renal responses to FGF23 in the setting of kidney disease.(11,22) Here, we established a role for the STAT1-CEBPB-NFKB1 cascade in adverse signaling outcomes of FGF23 during DKD. Although the STAT1 activation was present in vehicle-treated mice with DKD in the PT-S1 segment, a substantial STAT1 signal was found in several nephron segments upon FGF23 treatment, whereas STAT1 was not among the most active TFs in any segment of the healthy groups. Notably, CEBPB TF activity was strongly induced by FGF23 in DKD but was not restricted to the direct FGF23-responsive nephron segments where presence of co-receptor Klotho facilitates canonical FGF23-FGFR signaling. (28) Rather, the broad CEBPB TF activity along the nephron may be induced by other early signals, the identity of which remains elusive. Cytokines such as soluble TNF receptors 1 and 2 or C-reactive protein are induced by states of kidney disease and/or by FGF23 (11,22) and could initiate proinflammatory transcriptional programs along the nephron.

### Comparison with the literature

Agoro et al. recently dissected FGF23 effects along the nephron in healthy C57BL/6 mice.(10) We confirm some of their findings in our FGF23-treated healthy DBA/2J mice, including FGF23-driven *Egr1* expression and EGR1 regulon activity along the nephron. A discrepancy was that our single-nucleus study found FGF23-driven *Cyp24a1* expression in both healthy PT-S1 and DCT1 segment, whereas the single-cell study of Agoro et al. detected *Cyp24a1* in PT-S1/S2/S3 but not in DCT. Both acute and also chronically sustained FGF23 signaling can induce expression of pro-inflammatory TNF receptor superfamily members even in otherwise healthy kidney.(10,29) Agoro et al. also predicted NF-KB activity as downstream consequence of FGF23-driven TNF receptor superfamily members expression.(10) The present data establish an important upstream layer with the CEBPB regulon. While Cebpb is known as key TF mediating proinflammatory effects in the vasculature (24) and in several inflammatory diseases (30), its role is not limited to diabetes. We previously reported that in a kidney disease model of Adriamycin-induced nephrotoxicity, renal *Cebpb* transcripts were increased 24h after a single injection with recombinant FGF23.(11) This previous finding corroborates the present study’s main finding using an orthogonal experimental strategy, in addition to the context provided by the literature on inflammation.

### Strengths and weaknesses

Strengths of the present work include the systematic profiling of hormone FGF23 renal effects in DKD using large study with the latest Chromium 10X Gene Expression kit able to load nearly 20.000 nuclei per sample. Further, we used a uniform low-dose treatment with FGF23 to obtain near-physiological instead of supraphysiological signaling effects. This study also has some limitations: Agoro et al. Show that in PT, DCT and CNT, a maximal expression of *Tnfrsf12a* and *Tnfrsf1a* at 4h treatment (10). Tnfrsf12a had a particularly elevated chromatin accessibility (10). The present work had a different focus and assessed an earlier time point during initial FGF23 signaling at 1h only, but we scaled the sequencing to a much deeper level with 4 animals per group and a higher number of nuclei per sample. It will be important to assess longitudinal effects of FGF23 at the single-cell level also in diseased kidney, and to assess how a longer-term Cebpb inhibition affects both DKD and FGF23 signaling in DKD. Next, a limitation of this study is the male-specific analysis of the DKD model, as we failed to generate a robust disease induction in females using the chosen protocol. The sex-specific dosing was chosen based on the known relative protection from STZ-induced diabetes due to estrogen in females (12,13), but the failure of the elevated STZ dose to induce DKD thus precludes sex-specific data interpretation in the present study. Finally, a further limitation is the fact that STZ-induced DKD increased phosphaturia in experimental mice even without FGF23 treatments. A similar phosphaturia increase has previously been described in STZ-treated rats (31). Thus, true disease-specific differences in FGF23 effects on phosphaturia are difficult to interpret in the chosen model.

### Implications

In physiology, already minor changes in cell signaling due to altered co-receptor (e.g. Klotho) abundance, receptor or kinase tropism, signaling feedback mechanisms and thus signaling outcomes may compound drastically over time. The present study identified Cebpb as a potential therapeutic target for aiming to alleviate the adverse effects of excessive FGF23 in an experimental DKD model. A strategy of longer-term Cebpb inhibition in DKD will be the next logical step, in addition to extending the validity to additional disease models. As shown by others before, genetic variation in *Cebpb* in mice and humans revealed *Cebpb* as a risk allele for podocyte loss and thus DKD severity.(27) Thus, there is excellent rationale to follow up on the present data with a translational focus on addressing CKD complications.

### Conclusions

Overall, we combined a mouse model with experimental DKD and single-nucleus analyses to link FGF23 signaling with activity of the STAT1-CEBPB regulon, a known driver of inflammation that can be pharmacologically modified. Our study reveals Cebpb as a novel therapeutic target to alleviate the proinflammatory actions of elevated circulating concentrations of FGF23 in DKD.

## Data availability

Single-nucleus RNAseq dataset has been deposited to GEO with accession number: GSE283997. Bulk RNAseq data are in the process of being deposited to GEO. Code will become available on GitHub. Raw phenotype data are available on request.

## Acknowledgements

The authors would like to acknowledge the Single cell core Facility at Flemingsberg campus Karolinska (SICOF), supported by the Karolinska Infrastructure board, for their single cell sequencing services. Computational resources were provided by the National Academic Infrastructure for Supercomputing in Sweden (NAISS), funded by the Swedish Research Council.

## Author contributions

Conceptualization: MBM, HO

Methodology: MBM, CY, SS, MB PC

Software: CY, SS, PC

Validation: MBM, CY, SS, MB

Formal analysis: MBM, CY, SS, MB, PC

Investigation: MBM, CY, SS, MB, PC, HO

Resources: MBM, MB, PC, JP, HO

Data Curation: MBM, CY, SS, MB

Writing - Original Draft: MBM, CY, SS

Writing - Review & Editing: MBM, CY, SS, MB, JP, PC, HO

Visualization: MBM, CY, SS

Supervision: MBM, PC, HO

Project administration: MBM, MB, PC, HO

Funding acquisition: MBM, PC, HO

