## Supplemental Figures for "Single-nucleus analysis reveals renal STAT1/CEBPB signaling as a modifiable relay of proinflammatory FGF23 effects in diabetic kidney disease"

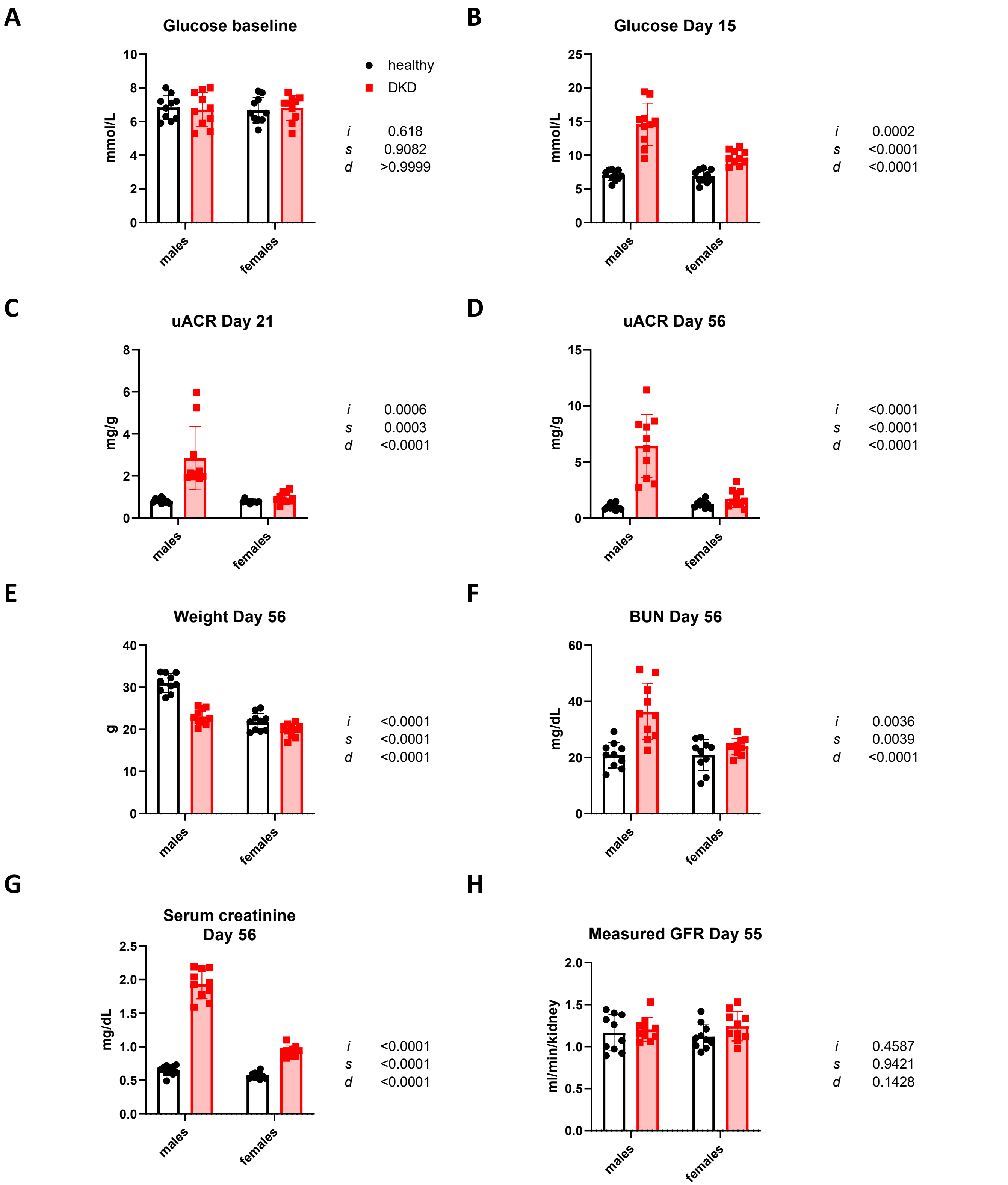


Supplemental Figure 1. Physiological analyses in mice of both sexes upon induction of diabetic kidney disease (DKD). Two-Way ANOVA shows p-values for contrasts of disease state (d), sex (s) and interaction (i). GFR, glomerular filtration rate. uACR, urinary albumin/creatinine ratio. N=5 per group.

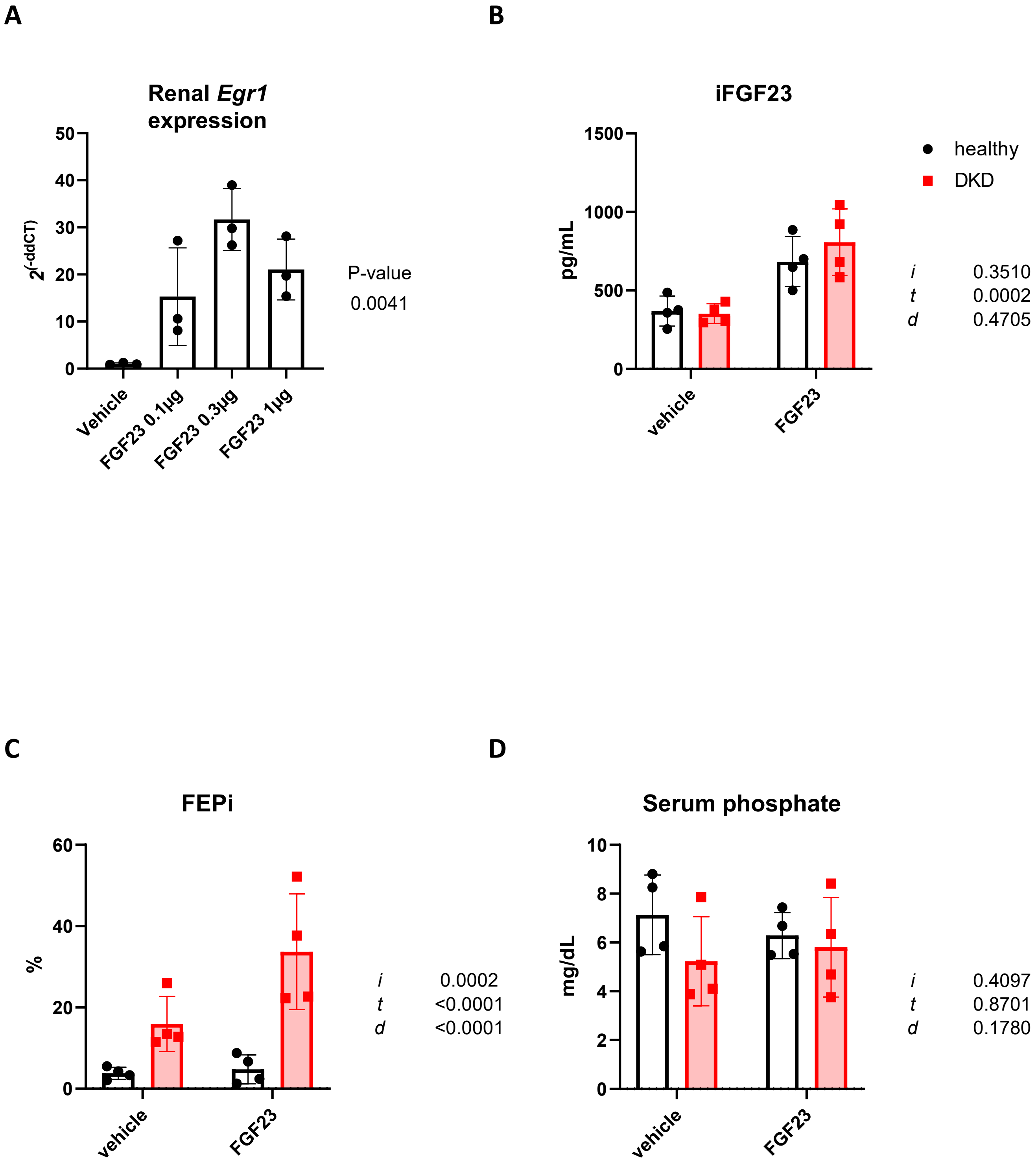


Supplemental Figure 2. Transcriptional analysis of FGF23 treatments at different doses (A). B-D show responses on serum FGF23 and phosphate metabolism inresponse to FGF23 in male mice. Analysis by ANOVA (A) or Two-Way ANOVA (B-D) with p-values shown for indicated contrasts of disease state (d), FGF23 hormone treatment (t) and interaction (i). FEPi, fractional excretion of phosphate. N=3 (A) or 4 per group (B-D).

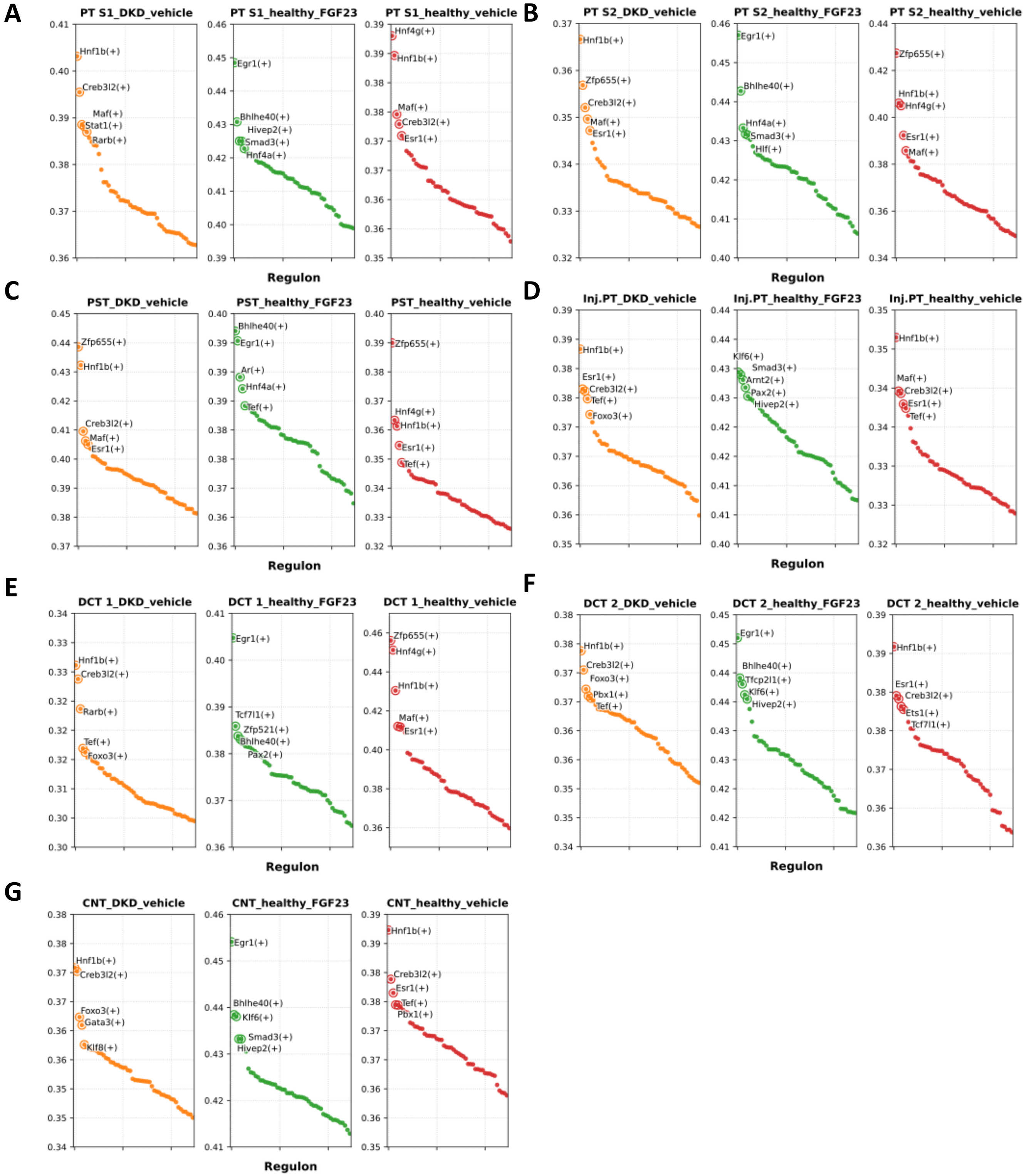


Supplemental Figure 3: Supporting figure to Figure 3. Regulon analysis by pySCENIC stratified by experimental group, in 4 proximal and 3 distal nephron segments. PT, proximal tubule. PST, proximal straight tubule. CNT, connecting tubule. DCT, distal convoluted tubule. S1-3, segment 1-3. N=4 mice per group. Y axis denotes regulon specificity score.

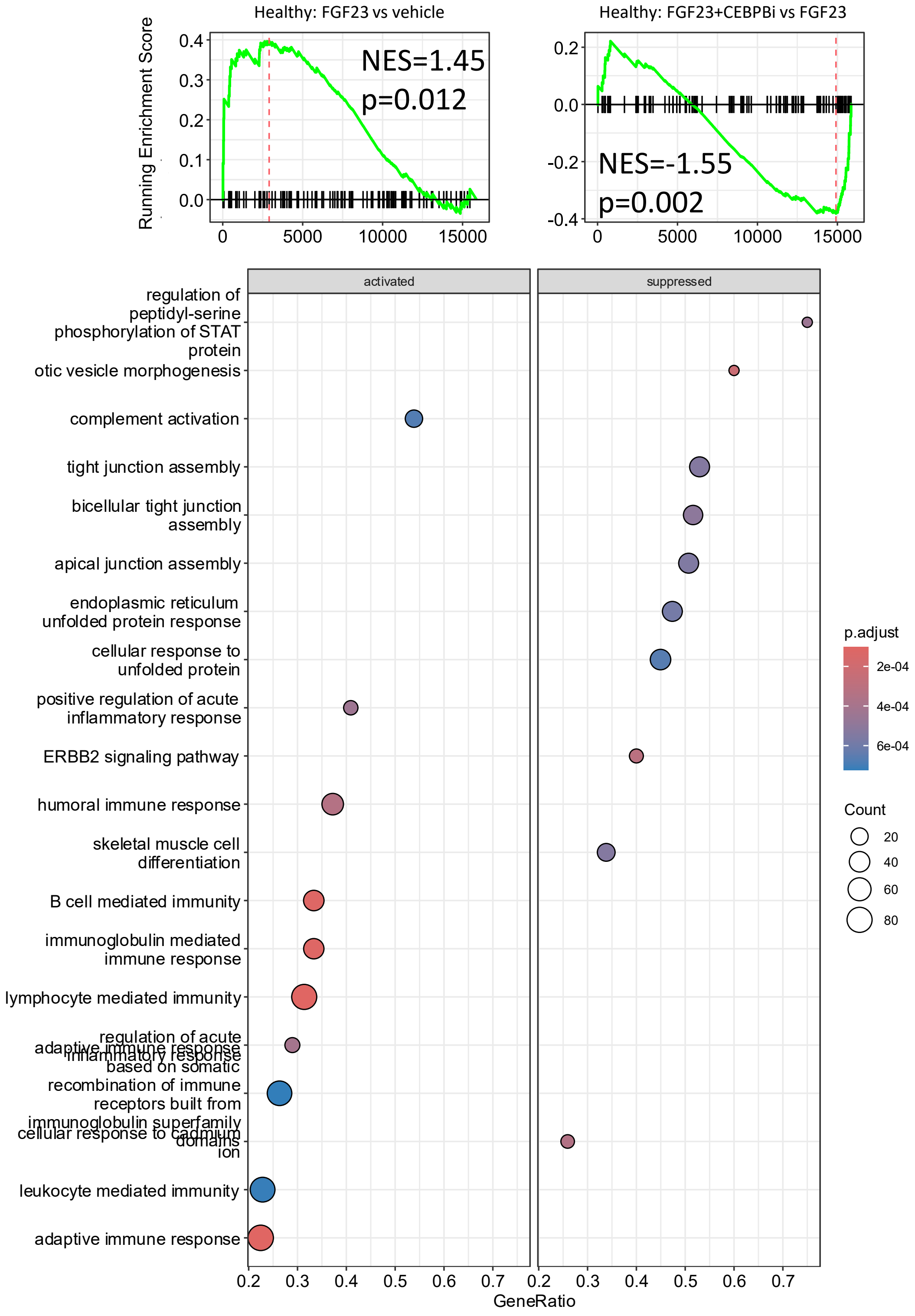


Supplemental Figure 4: Effect of FGF23 or CEBPB inhibitor ST-101 on CEBP target gene set (top) and on overall top enriched gene sets (bottom) of FGF23 signaling in precision-cut kidney slices from healthy mice. Contrasts as indicated (top), and lower panel: Gene set enrichment of FGF23 + ST-101 vs. FGF23, paired contrast. N=3 miced.

**CEBPBi effect on TF activity during FGF23 signaling in healthy kidney**

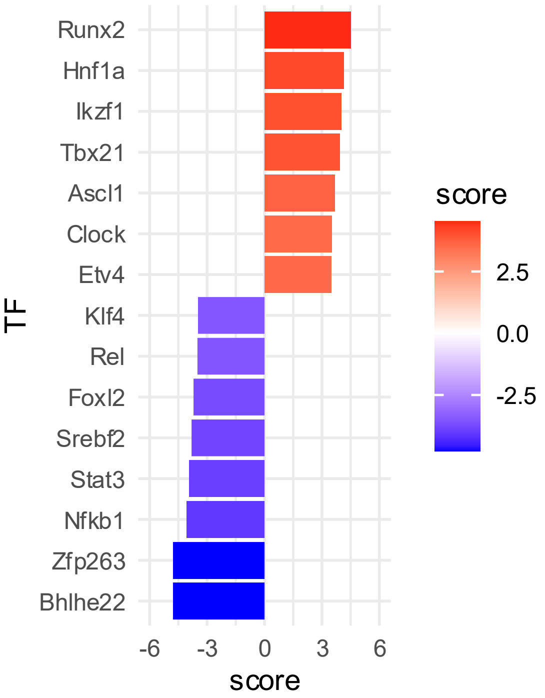


Supplemental Figure 5: Effect of CEBPB inhibitor (CEBPBi) ST-101 effects on transcriptome-inferred transcription factor (TF) activity during FGF23 signaling in precision-cut kidney slices from healthy mice, using t statistic of paired limma contrast of FGF23 + ST-101 vs. FGF23. N=3 mice, top 15 TF displayed.
